# Differentiating Bayesian model updating and model revision based on their prediction error dynamics

**DOI:** 10.1101/2022.06.15.496278

**Authors:** Danaja Rutar, Olympia Colizoli, Luc Selen, Lukas Spieß, Johan Kwisthout, Sabine Hunnius

## Abstract

Within predictive processing learning is construed as Bayesian model updating with the degree of certainty for different *existing* hypotheses changing in light of new evidence. Bayesian model updating, however, cannot explain how *new* hypotheses are added to a model. Model revision, unlike model updating, makes structural changes to a generative model by altering its causal connections or adding or removing hypotheses. Whilst model updating and model revision have recently been formally differentiated, they have not been empirically distinguished. The aim of this research was to empirically differentiate between model updating and revision on the basis of how they affect prediction errors and predictions over time. To study this, participants took part in a within-subject computer-based learning experiment with two phases: updating and revision. In the updating phase, participants had to predict the relationship between cues and target stimuli and in the revision phase, they had to correctly predict a change in the said relationship. Based on previous research, phasic pupil dilation was taken as a proxy for prediction error. During model updating, we expected that the prediction errors over trials would be gradually decreasing as a reflection of the continuous integration of new evidence. During model revision, in contrast, prediction errors over trials were expected to show an abrupt decrease following the successful integration of a new hypothesis within the existing model. The opposite results were expected for predictions. Our results show that the learning dynamics as reflected in pupil and accuracy data are indeed qualitatively different between the revision and the updating phase, however in the opposite direction as expected. Participants were learning more gradually in the revision phase compared to the updating phase. This could imply that participants first built multiple models from scratch in the updating phase and updated them in the revision phase.

## Introduction

Imagine you recently moved from one desert city in Australia to another. In your old neighbourhood all front yards were brown due to severe drought. However, in your new neighbourhood, to your surprise, all the front yards appear to be green. You wonder how this is possible since both cities have the same climate. You come up with potential reasons for this and decide that probably the new neighbourhood has just experienced a period of heavy rain. It is not until a week later that you see your neighbour changing their lawn for a new one. Now, you realise that all the front yards in this town have artificial green lawns. At that moment you add a new hypothesis to your model that you did not possess before.

What challenge does the example above pose? We make sense of the world through models and use them for generating hypotheses about what the causes of sensory evidence are (Clark, 2016; Kwisthout et al., 2017). When a model fails to adequately predict the sensory evidence, it needs to be adjusted. In the case above, a new hypothesis needs to be added to account for the new observation. How does this happen? What mechanism allows for the integration of new hypotheses into an existing model? This question has been one of the central questions in cognitive science, but it still remains unanswered.

Here, we address this long-standing question within a predictive-processing framework, a popular and influential theoretical framework in computational cognitive neuroscience (e.g., Friston et al., 2009; Friston et al., 2016; Friston et al., 2017; Clark, 2013; 2016; Hohwy, 2013). The predictive-processing framework aims to become a unifying framework for understanding the entirety of human cognition and behaviour from visual processing (Rao & Ballard, 1999; Edwards et al., 2017; Petro & Muckli, 2016) to mentalizing (Kilner et al., 2007; Koster-Hale & Saxe, 2013). According to the theory, the brain embodies a hierarchical generative model that “aim[s] to capture the statistical structure of some set of observed inputs by tracking (one might say, by schematically recapitulating) the causal matrix responsible for that very structure” (Clark, 2013, p. 2). Based on this hierarchical model, the brain generates top-down predictions, which are compared to the incoming sensory signals. The difference between the predicted and the actual sensory input, i.e., the prediction error, is computed. From a predictive-processing perspective, minimising reducible prediction error is the primary goal of computations in the brain and occurs mainly as a result of learning (Friston et al., 2016; Smith et al., 2020; Fitzgerald, 2015; Clark, 2016; Zenon, 2019).

Existing theories of learning in predictive-processing make use of Bayesian models of cognition. As a natural consequence of using the Bayesian framework, most research on learning has focused on one specific form of learning, Bayesian model updating, which explains how model hypotheses are *updated* in the light of new evidence using Bayes’ rule (da Costa et al., 2020; Friston et al., 2016; Smith et al., 2020). In other words, every time new sensory evidence comes in, hypotheses in the learner’s model gain and lose probability (Kruschke, 2014; Perfors et al., 2011; Gopnik & Tenenbaum, 2007; Tenenbaum et al., 2006). Learning accounts within the Bayesian framework are well suited for explaining updating of hypotheses and selection among these existing hypotheses. However, they assume that the entire set of relevant hypotheses and the mechanism for comparing hypotheses are predefined. Learning under this view is reduced to *choosing* among existing hypotheses, whereas the process of *generating* novel hypotheses is neglected (Perfors, 2012). Unless one assumes that learners are equipped from the start with the complete set of hypotheses that they will ever consider, one needs to account for how novel hypotheses are *added* to an existing model (Gentner & Hoyos, 2017; Schulz, 2012; Christie & Gentner, 2010). Bayesian updating, however, does not account for this creative aspect of learning (Perfors, 2012; Gentner & Hoyos, 2017; Schulz, 2012; Xu, 2019; Christie & Gentner, 2010; Kwisthout et al., 2017). Kwisthout and colleagues (2017) proposed that learning of novel hypotheses should be computationally differentiated from Bayesian model updating and coined the term “model revision” for this alternative learning mechanism. Model revision, unlike model updating, changes the structure of a generative model by altering its causal connections or by adding and removing hypotheses (Kwisthout et al., 2017). Similarly, Heald et al. (2022) have recently presented a theory for sensorimotor learning, called contextual inference, that differentiates between the adaptation of behaviour based on updating of existing and creation of new motor memories and adaptation due to changes in the relative weighting of these motor memories.

Building on the theoretical argument that model updating and model revision are computationally distinct, the aim of this study was to investigate whether these two learning mechanisms can be empirically distinguished. To investigate this, we created an experiment with two phases: an updating and a revision phase. In the updating phase, participants had to learn the relationship between cues and a target stimulus. We induced the need for model updating through a change in this relationship and examined whether the two experimental phases would lead to two distinct patterns of learning. More specifically, we hypothesised that the change in prediction error over time would follow distinct trajectories in the model updating and model revision phases. In the model updating phase, the prediction error should *gradually* decrease due to gradual updating of the probability distribution of existing model hypotheses. In the model revision phase, prediction error should *abruptly* decrease once a new model hypothesis has been added to the model (see section Hypotheses for a more detailed description).

Whilst prediction error cannot be measured directly, it has been assessed indirectly, for instance using EEG (Sambrook et al., 2018; Brydevall et al., 2018; Maier et al, 2019) or pupillometry (Preuschoff et al., 2011; Nassar et al., 2013; Browning et al., 2015; O’Reilly et al., 2013; Colizoli et al., 2018; Lawson et al., 2017). In this study, we examined pupil dilation as a proxy for so-called cognitive prediction error (den Ouden et al., 2012).

We hypothesised that prediction-error dynamics resulting from model updating and model revision should also be reflected in participants’ predictions of the upcoming target stimuli. Prediction error minimisation is the result of learning more adequate models of the world (Clark, 2013; 2016; Hohwy, 2013), reflected in the fact that predictions gradually become more correct over time. We thus expected that in the model updating phase, predictions would only gradually become better, whereas in the model revision phase, predictions should show a more instantaneous change from incorrect to correct. Our initial idea was to study model updating and model revision based on their *prediction error* minimisation dynamics. However, the results we obtained were unexpected and, therefore, in addition to investigating prediction error minimisation dynamics, we also performed an exploratory behavioural analysis focusing on *predictions* based on button presses.

## Method

### Participants

Participants were recruited using Radboud University’s online recruitment system. The only restriction for participation was a minimum age of 16 years. Thirty-three healthy adults with normal or corrected-to-normal vision participated in our study. One participant was excluded for not following the instructions properly. Two participants were excluded due to equipment malfunction. The final sample consisted of 30 participants (9 males, 21 females, mean age = 23.3, range: 19 – 42 years). The Ethical Committee of the Social Sciences Faculty at Radboud University approved the study, and all participants gave written informed consent. Participants received 15 euros for participating in the experiment.

### Task and procedure

Participants carried out a computer-based two-alternative forced-choice (2AFC) task on the *expected* orientation (left vs. right) of Gabor patches (Fig 1A). All participants took part in two consecutive phases of the task, first the updating and then the revision phase. The entire experiment took 1.5 hours and had three breaks (two short breaks halfway each phase and one longer break between the two phases). After each break, recalibration of the eye-tracker took place (see below). At the beginning of the experiment, participants were told that they would be presented with auditory and visual cues. The instructions were to use the cues to predict the orientation of the target stimulus in each trial. Before the start of the experiment, participants were presented with an example trial. They indicated their prediction by pressing either the right or left button on a button box for left or right orientation, respectively. Participants were instructed to press the button as soon as they thought they knew which target orientation would appear. At the end of the updating phase, participants were told that the cue-target rule would change (i.e., the model revision phase) but not what the change would be. In the revision phase, participants were similarly instructed to continue to use the cues to predict the orientation of the target stimulus. The ‘model revision’ was a reversal of the cue-target contingencies in the revision phase only for the trials which contained an auditory tone. Each phase (updating and revision) of the experiment consisted of 200 trials (400 trials in total).

**Figure 1.**
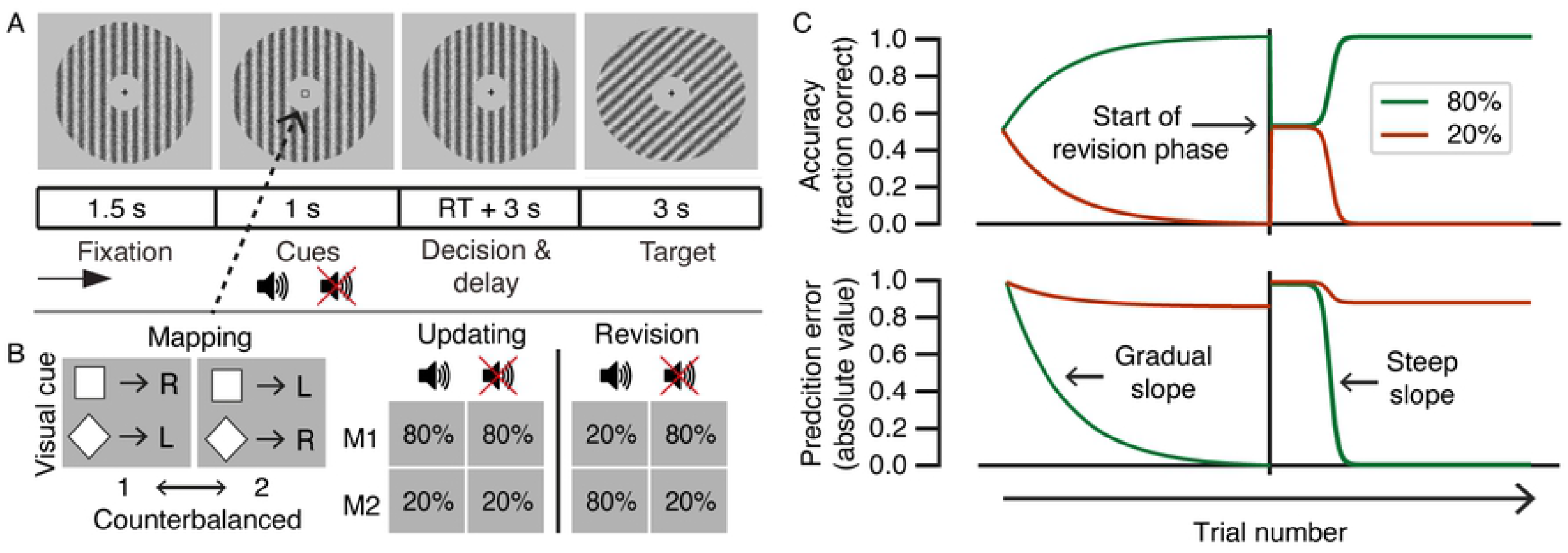
Experimental design and hypotheses. (**A**) Behavioural task. Participants performed a 2AFC task on the expected orientation (left/right) of upcoming Gabor patches while pupil dilation was recorded. Each trial consisted of a fixation period, a cue period, a response window followed by a delay period, and finally a target period. The decision interval ranged from onset of the cue to the participant’s response. The target interval ranged from target onset into the subsequent inter-trial interval (3 s). The target served as feedback on the accuracy of participants’ predictions in the decision interval. (**B**) The participants had to learn cue-target contingencies in order to accurately predict the orientation of the upcoming Gabor patch (target). Mapping 1 (M1) was defined as the visual cue-target pairs that occurred in 80% of trials in the updating phase; Mapping 2 (M2) was defined as the visual cue-target pairs that occurred in 20% of trials in the updating phase. Mappings were counterbalanced between participants. At the start of the revision phase, the frequencies (80% vs. 20%) of the cue-target mappings were reversed for trials containing the auditory tone cue only. (**C**) Main hypotheses for the dynamics of prediction errors and accuracy over the course of the 2AFC task. The first 200 trials of the task represent the updating phase in which a gradual increase in accuracy and a gradual decrease in the absolute value of prediction errors were expected (represented by exponential curves). Within the revision phase (the last 200 trials), an abrupt increase in accuracy and an abrupt decrease in prediction errors were expected (represented by sigmoidal curves).

All participants were seated approximately 50 cm from the screen with their chin in a chin rest. Stimuli were isoluminant and the environmental illumination was the same for all participants. Stimuli were presented on a computer screen with the spatial resolution of 1920 × 1080 pixels. One trial lasted approximately nine seconds. Each trial consisted of a fixation period, a cue period, a response window followed by a delay period, and a target period. For all periods, except for the target period, a vertically oriented Gabor patch was presented. The target stimulus was a Gabor patch oriented to the left or to the right, with a special frequency of 0.033 and opacity 0.5. Each trial started with a fixation cross on the vertical Gabor patch, which was shown on the screen for 1500 ms. Afterward, the visual cue, which was either a square or a diamond, was presented in the middle of the screen for 1000 ms. In 50% of the trials the visual cue was paired with an auditory cue (i.e., tone “C”), which was presented for 300 ms. In the other 50% of the trials, the auditory cue was absent. During the subsequent interval, the fixation cross and the vertically oriented Gabor patch were presented, and participants were asked to indicate their prediction about the upcoming orientation of the target by pressing a button. There was no maximum response window. After a button had been pressed, a delay period started with the vertical Gabor patch on the screen for additional 3000 ms. After the delay period, the target stimulus (the Gabor patch tilted either left or right) was shown for 3000 ms. The durations of the response period and target window were chosen in order to avoid contamination of the pupil dilation response to a previous event. As our dependent measure was the pupil dilation response to the target stimulus, the response window was sufficiently long to ensure that the pupil response would not be contaminated by the motion response of the button press (Colizoli et al., 2018). It is important to note that the target Gabor patch served as trial-by-trial feedback on the accuracy of participants’ cue-target predictions. The prediction error in this context is therefore the difference between the expected orientation of the target and the target’s actual orientation.

The design contained probabilistic cue-target mappings to introduce uncertainty in the predictions, thus simulating uncertainty that is inherent to perception in the real world. The visual and auditory cues predicted whether the target Gabor patch was tilted to the right or to the left with either an 80% or 20% probability (Fig 1B). In the updating phase, only the visual cue had predictive value for learning the cue-target probabilities (i.e., the auditory tone did not predict the target). In the revision phase, the visual cue-target mappings were reversed only for trials in which an auditory cue was presented. Cue-target mappings were counterbalanced between participants. Note that we define mapping 1 (M1) to correspond to the 80% visual cue-target pairs in the updating phase and mapping 2 (M2) to correspond to the 20% visual cue-target pairs in the updating phase.

### Hypotheses

#### Main hypotheses

We hypothesised that in the updating phase, participants would be gradually learning the probabilistic relationship between the cues and the Gabor patch orientation (target). New sensory evidence, in this case the target-stimulus presentation, should deviate less from expectations as evidence is accumulated over trials and thus generate smaller prediction errors for the high frequency trials (i.e., their absolute values). We thus expected that in the updating phase, the prediction errors would show a *gradual* decrease over time, illustrated by an exponential curve. At the beginning of the revision phase, participants were instructed that something would change during this revision phase. We expected that participants would discover the new rule by integrating the now meaningful tone into their predictive models. The new hypothesis should account for observations that were previously generating high prediction errors. After an initial increase (compared to the ending point in the updating phase) in prediction errors in the revision phase, an integration of a new hypothesis should lead to an *abrupt* decrease in prediction errors (i.e., an “aha” moment), illustrated by sigmoidal curves. Fig 1C (bottom) illustrates the expected prediction error during the 2AFC task. We expected a similar dynamic for the tone and no tone trials in the updating and revision phase, however, it could be that the learning curves for these trial types differed during revision due to the change in contingency. Therefore, we investigated the tone and no-tone trials separately.

#### Exploratory hypotheses

While prediction errors reflect the deviation between expectations and feedback presentation, the behavioural responses (i.e., button presses) are supposed to reflect the *predictions* themselves. The experimental manipulation should affect accuracy data in a similar way to prediction errors but in the opposite direction. We hypothesised that in the updating phase, the accuracy of predictions should increase gradually and then plateau, while in the revision phase, accuracy should initially decrease then abruptly plateau again as soon as the change in the cue-target contingency is learned.

#### Estimating prediction errors from pupil dilation

Under constant luminance conditions, pupils dilate more when an error is made as compared with a correct response (e.g., Braem et al., 2015). Recently, it was shown that pupil dilation furthermore reflected an interaction between response accuracy and task difficulty in a 2AFC motion-discrimination task with reward-coupled feedback, indicating its potential for investigating reward prediction errors (Colizoli et al., 2018). In this previous work, the effect of response accuracy (errors vs. correct trials) in the feedback-locked pupil dilation was larger for hard trials as compared with easy trials on average. Following these results, we investigated whether target-locked pupil dilation could be used as a measure of prediction errors in the current study.

### Data acquisition and analyses

#### Data acquisition and pre-processing

Changes in pupil dilation were recorded using an SMI RED500 eye-tracker (SensoMotoric Instruments, Teltow/Berlin, Germany) with a sampling rate of 500 Hz. We analysed the pupil dilation data of the right eye for each participant. The timing of blinks and saccade events was not saved in the output of the eye-tracker; therefore, we did not attempt to categorize separate blink and saccade events for pre-processing purposes. Pre-processing was applied to the entire pupil dilation time series of each participant and consisted of: i) interpolation around missing samples (0.15 s before and after each missing period), ii) interpolation around blinks or saccade events based on spikes in the temporal derivative of the pupil time series (0.15 s before and after each blink or saccade period), iii) band-pass filtering (third-order Butterworth, passband: 0.01–6 Hz), iv) removing responses to nuisance events using multiple linear regression (missing periods and blink or saccade events were all categorized together as a single ‘nuisance’ event type; responses were estimated by deconvolution, [Knapen et al., 2016]), and v) the residuals of the nuisance regression were transformed to percent signal change with respect to the temporal mean of the time series.

For each trial, intervals corresponding to the onset of the cue were extracted from each participant’s pupil dilation time series (cue-locked and target-locked, respectively). The cue-locked and target-locked intervals were baseline-corrected separately for each trial. The baseline pupil was defined as the mean pupil in the time window −0.5 to 0 seconds with respect to the cue or target for the cue-locked and target-locked intervals, respectively. The cue-locked pupil response was analysed for data quality purposes, while the target-locked pupil response was the main dependent variable of interest.

The temporal window of interest was independently defined as 1 to 2 seconds after the target onset based on the pupil’s canonical impulse response function (Knapen et al., 2016). For each trial, a single value for the target-locked pupil response was computed as the mean pupil dilation within this temporal window of interest. The event-locked and baseline-corrected scalar value representing pupil dilation on each trial is referred to as the phasic pupil response in the literature. Note that the pre-trial baseline pupil dilation is expected to reflect tonic fluctuations in arousal in contrast to the task-evoked (or phasic) pupil responses investigated in the current study (Murphy, O’Connell et al., 2014; Murphy, Vandekerckhove et al., 2014).

Trials were excluded if the reaction time (RT) was three standard deviations above the participant’s mean RT or lower than 100 ms (minimal time needed for the preparation of a motor response; Dahan et al., 2019; Siegelman et al., 2019).

#### Confirmatory analyses

##### Behaviour

We expected that, on average, participants would learn the cue-target contingencies in both phases of the task, which would be reflected in higher accuracy and faster responses for high frequency trials as compared with low frequency trials. The effect of the visual cue-target mapping condition was expected to interact with the auditory cue condition and task phase, reflecting the reversal of the visual cue-target contingencies in the revision phase during the tone trials only. These hypotheses were tested in two 3-way repeated measures ANOVAs, separately for accuracy (as percentage of correct trials) and RT with factors: cue-target mapping (M1 vs. M2), auditory cue (tone vs. no tone), and phase (updating vs. revision).

##### Evoked pupil response

The analysis of the evoked pupil response due to the cue presentation served as a quality-control check on the goodness of the data acquisition. The visual and/or auditory cues indicated to the participants that they should make a button press based on their choice. A button press in the response phase was expected to evoke a canonical pupil impulse response which should be reflected in the mean cue-locked pupil response. In addition, we expected to see larger pupil dilation on average during tone trials as compared with no tone trials in the cue-locked pupil response, as auditory cues are known to be arousing (e.g., Liao et al., 2016). The cue-locked effect of tone was expected to return to baseline before the target was presented on screen. We expected to see larger pupil dilation on average during errors as compared with correct trials in the evoked target-locked pupil response.

##### Phasic pupil response

Following from the hypothesised pattern of behavioural results, we expected to see an interaction between cue-target mapping, auditory cue, and task phase in the target-locked phasic pupil response in the average phasic pupil response (this was tested with a 3-way repeated measures ANOVA as described above for behaviour). In the updating phase, the average pupil dilation for the M2 mapping was expected to be larger as compared with the M1 mapping, because low frequency trials (M2 in updating) should contain more errors overall. The direction of the mapping effect was expected to reverse in the revision phase for tone trials only.

##### Curve fits on target-locked pupil data

To test our main hypothesis concerning prediction error dynamics across the updating and revision phases, we assessed whether the time course of the target-locked pupil dilation showed the difference in the range parameter, σ, of the sigmoid curve fits. The target-locked pupil dilation is taken as a proxy of prediction errors within two phases of the experiment. The logic is that sensory evidence for the cue-target continencies is accumulated as the task progresses over time, and learning should be evident in a reduction in prediction error magnitude as a function of trial order. The sigmoid function used for the curve fits on target-locked pupil responses is given in Equation 1.

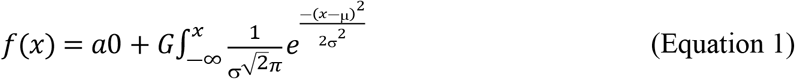

We fit four parameters, μ, σ, a0 and G, of the above sigmoid function to the target-locked pupil dilation across the high-frequency trials (80%) for the tone condition only, comparing the updating and revision phases of the experiment. We differentiated between the high-frequency as compared with low-frequency trials, in order to fit only a single direction of the (expected) signed prediction errors (Colizoli et al., 2018; see den Ouden et al., 2012). The starting point of the curve is reflected in the a0 parameter. The inflection point of the curve is reflected in the μ parameter. The gain parameter, G, allowed for negative scaling of the curves given the nature of the pupil signal as dependent variable (i.e., percent signal change). The range parameter, σ, reflects the range over where the curve rises. A larger σ parameter is associated with a larger range over which the transition (i.e., from f(x) = a0 to f(x) = 1) takes place. We placed linear constraints on the curve fits so that σ could not exceed three times the value of μ. We bound both μ and σ so they could not exceed the number of trials in each phase of the experiment (range between 1 and 200 trials). The parameters were furthermore not constrained or bounded. The parameters were determined by minimising the ordinary least squares cost function.

Our main hypothesis was that learning would extend across more trials in the updating phase as compared with in the revision phase, reflecting the difference between model updating and model revision processes. Therefore, we expected that the σ would be higher for the high frequency tone trials in the updating phase as compared with the high frequency tone trials in the revision phase (note that the cue-target mappings were flipped for tone trials in the revision phase; see Fig 1C). We furthermore expected the sign of the G to be negative in both phases, indicating a decreasing trend. A larger value of σ together with a negative gain parameter, G, thus reflects a more gradual reduction of prediction errors across trials. This analysis enables us to examine whether pupil dilation depended on the experimental phase and if it scaled with our hypotheses in Figure 1C. Descriptive statistics for all free parameters are presented in Supplementary Table 1.

Finally, in the Supplementary Materials we describe the prediction errors as the 4-way interaction between accuracy, cue-target mapping, auditory cue, and phase in the target-locked pupil dilation.

### Exploratory analyses

#### Psychometric curve fits on accuracy data

To test the hypothesis concerning the dynamics of the target-orientation predictions, we assessed whether the range parameter, σ, of the psychometric curve fits differed between the updating and revision phases. In a psychometric curve, accuracy is plotted against signal intensity (May & Solomon, 2013). We explored the resulting psychometric curves when the number of trials completed over time was taken as a proxy for signal intensity. The sigmoid function used for the psychometric curve fits is given in Equation 2.

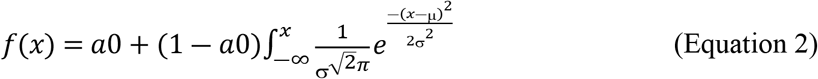

We fit three parameters, μ, σ, and a0, to the individual participant’s response accuracy across all trials for the tone condition only (i.e., the frequency conditions were not differentiated), separately for the updating and revision phases of the experiment. We placed linear constraints on the curve fits so that σ could not exceed three times the value of μ. We bound both μ and σ so they could not exceed the number of trials in each phase of the experiment (range between 1 and 200 trials). The starting point, a0, was bound between 0 and 1. The parameters were determined by minimising the negative log-likelihood cost function.

If our hypotheses about the difference between model revision and model updating are correct, then σ should be higher for the tone trials in the updating phase as compared with the tone trials in the revision phase. Descriptive statistics for all free parameters are presented in Supplementary Table 1.

### Software

The prediction task was administered with Psychopy (version 1.81; Psychopy, 2018). The behavioural and pupil data were processed with custom software using Python (version 3.6; Python Software Foundation, 2016). The evoked pupil responses were statistically assessed with a cluster-level permutation test as part of the MNE-Python package (Gramfort et al., 2013). Repeated measures ANOVAs and Bayesian tests were carried out in JASP (version 0.13.1; JASP Team, 2020). All data and code are publicly available (https://github.com/colizoli/learning_environmental_contingencies).

## Results

The current study aimed to compare prediction-error dynamics, quantified in terms of pupil dilation, during model updating and model revision. As predictions become more accurate over time, due to learning more adequate models of the world, prediction errors are minimised (Clark, 2013; 2016; Hohwy, 2013). Model revision, unlike model updating, changes the structure of a generative model by altering its causal connections or by adding and removing hypotheses (Kwisthout et al., 2017). To test these hypotheses, participants performed a 2AFC task on the *expected* orientation (left vs. right) of an upcoming Gabor patch (target) task while pupil dilation was recorded (Fig 1A). The participants had to learn cue-target contingencies in order to accurately predict the orientation of the target. Cue-target contingencies changed at the start of the revision phase in the following way: the cue-target mapping was reversed but only for trials containing the auditory tone cue. The target served as feedback on the accuracy of participants’ predictions in the decision interval.

### Confirmatory analyses

We first evaluated whether participants performed the 2AFC task as expected. Main effects and interactions between the visual cue-target mapping condition (M1 vs. M2), task phase (updating vs. revision), and the presence of the auditory cue (tone vs. no tone) were assessed in three independent 3-way repeated measures ANOVAs on the dependent variables of mean accuracy (Fig 2A), RT (Fig 2B), and target-locked pupil dilation (Fig 2C) (see Fig 2 for the analysis of evoked pupil responses). The ANOVA results are presented in Table 1, and relevant post hoc comparisons are presented in Fig 2. At the mean group level, a significant 3-way interaction was obtained between visual cue-target mapping condition, auditory cue condition, and task phase for accuracy and target-locked pupil response, but not for RT. Participants accurately predicted the cue-target contingencies in both phases of the experiment, illustrated by the main effect of visual cue-target mapping condition in the updating phase and the mapping reversal in the revision phase for tone trials only (Fig 2A). As expected, the target-locked pupil response was larger for the M2 trials as compared with the M1 trials during the updating phase, and the presence of the auditory cue reversed the direction of the target-locked pupil response in the revision phase for the tone trials (Fig 2C). Supplementary Fig 1 illustrates how the behaviour and target-locked pupil dilation changed as a function of time (25 trials per bin).

**Figure 2.**
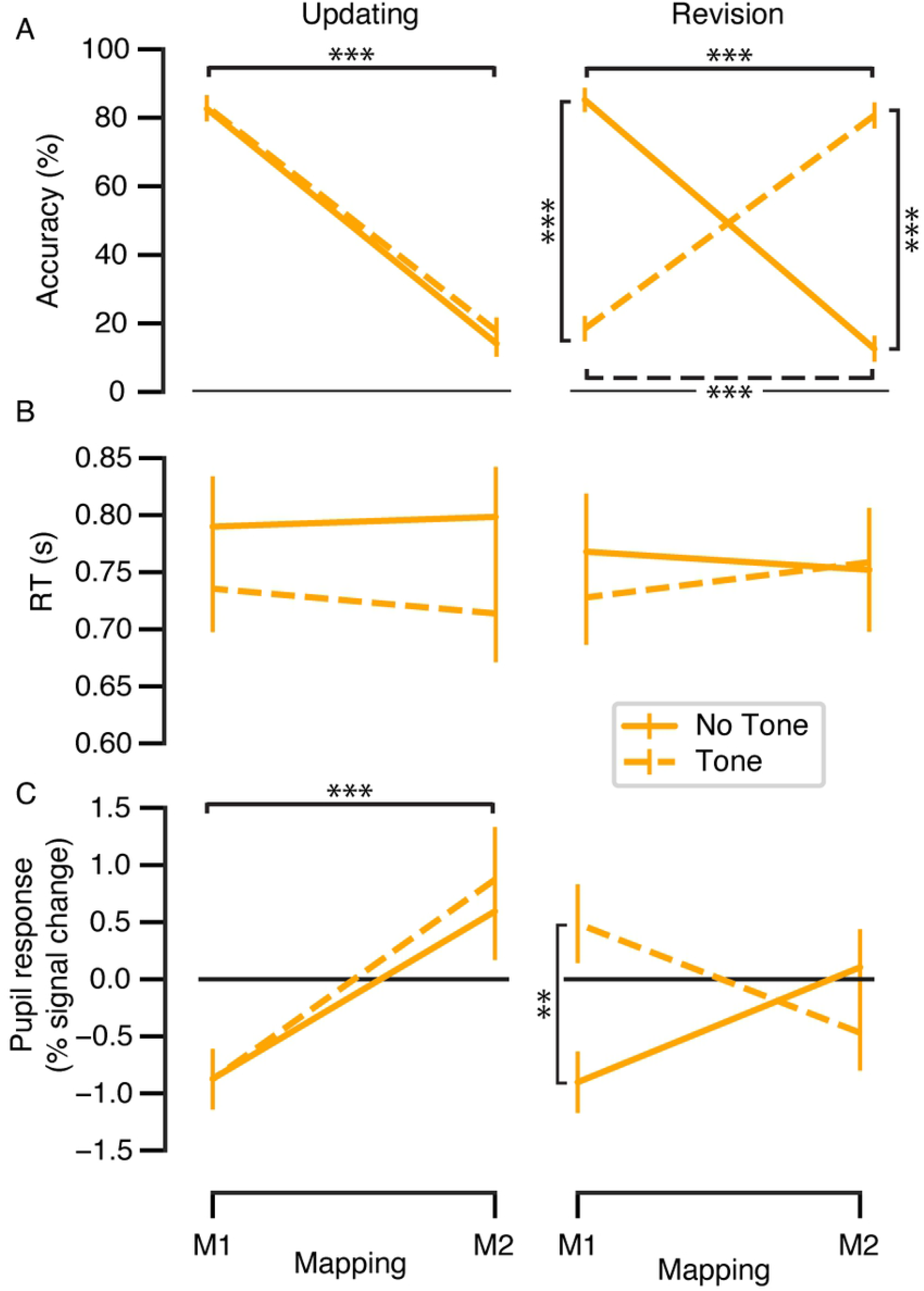
Cue-target prediction task results. (**A**) Prediction accuracy, (**B**) mean RT, and (**C**) target-locked pupil dilation as a function of visual cue-target mapping condition (M1 vs. M2), the presence of the auditory cue (tone vs. no tone), and task phase (updating vs. revision). Results of the 3-way repeated measures ANOVAs are given in Table 1. Significance refers to post hoc t-tests: \*\**p* < .01, \*\*\**p* < .001. Error bars, s.e.m. (*N* = 30). Note that the frequencies of the visual cue-target mappings change in the revision phase for the tone trials.

**Table 1.** Results of the 3-way repeated measures ANOVAs on accuracy, RT, and target-locked pupil response. Factors of interest were cue-target mapping condition (levels: M1 vs. M2), auditory cue (levels: tone vs. no tone), and task phase (levels: updating vs. revision). Accuracy data were percentage of correct trials; RT was analysed in seconds. \**p* < .05, \*\**p* < .01, \*\*\**p* < .001

| Effect | Accuracy |  |  | RT |  |  | Pupil response |  |  |
| --- | --- | --- | --- | --- | --- | --- | --- | --- | --- |
| | *F*(1,29) | *p* | $\eta^2_G$ | *F*(1,29) | *p* | $\eta^2_G$ | *F*(1,29) | *p* | $\eta^2_G$ |
| Mapping | 301.22 | < .001*** | 0.68 | < .01 | 0.964 | < .01 | 21.17 | < .001*** | 0.05 |
| Auditory cue | 6.69 | 0.015* | < .01 | 4.91 | 0.035* | 0.01 | 3.67 | 0.065 | 0.01 |
| Phase | 0.01 | 0.939 | < .01 | 0.14 | 0.711 | < .01 | 0.69 | 0.414 | < .01 |
| Mapping * Auditory cue | 226.34 | < .001*** | 0.65 | 0.16 | 0.694 | < .01 | 6.79 | 0.014* | 0.01 |
| Mapping * Phase | 201.03 | < .001*** | 0.61 | 0.39 | 0.535 | < .01 | 20.10 | < .001*** | 0.04 |
| Auditory cue * Phase | 0.40 | 0.533 | < .01 | 3.24 | 0.082 | < .01 | 0.57 | 0.458 | < .01 |
| Mapping * Auditory cue * Phase | 205.49 | < .001*** | 0.64 | 3.52 | 0.071 | < .01 | 9.71 | 0.004** | 0.02 |

The data quality of the pupil dilation measures was assessed with several checks based on previous literature before testing our hypotheses about the dynamics of prediction errors across the experimental phases. First, evoked pupil dilation was present in response to the (visual and auditory) cue onsets as expected in the decision phase, here reflecting both decision preparation as well as the upcoming motor output in the form of a button press (Colizoli et al., 2018; Urai et al., 2017) (Fig 3A). Furthermore, the temporal window (1 to 2 s) chosen for the phasic pupil analysis contained the peak of the group-level cue-locked evoked response as expected from the pupil’s canonical impulse response function (Fig 3A, grey box). Second, as expected, errors resulted in larger pupil dilation as compared with correct trials following the target presentation, and this accuracy effect was significant within the temporal window of interest (Fig 3B, grey box). Third, the presence of the auditory cue during the decision interval (cue-locked) was associated with larger pupil dilation as compared with the absence of the tone (Fig 3C), likely reflecting a difference in phasic arousal state during tone trials (Zekveld et al., 2018). Importantly, the (unwanted) arousal effect related to the auditory tone was no longer present by the time the target was presented for the participants (Fig 3D). We note that all further pupil analyses used the (phasic) target-locked pupil dilation, averaged within the temporal window of interest (Fig 3B, D, grey boxes), as the dependent variable. In sum, the pupil data fit all the basic confirmatory hypotheses based on previous literature.

**Figure 3.**
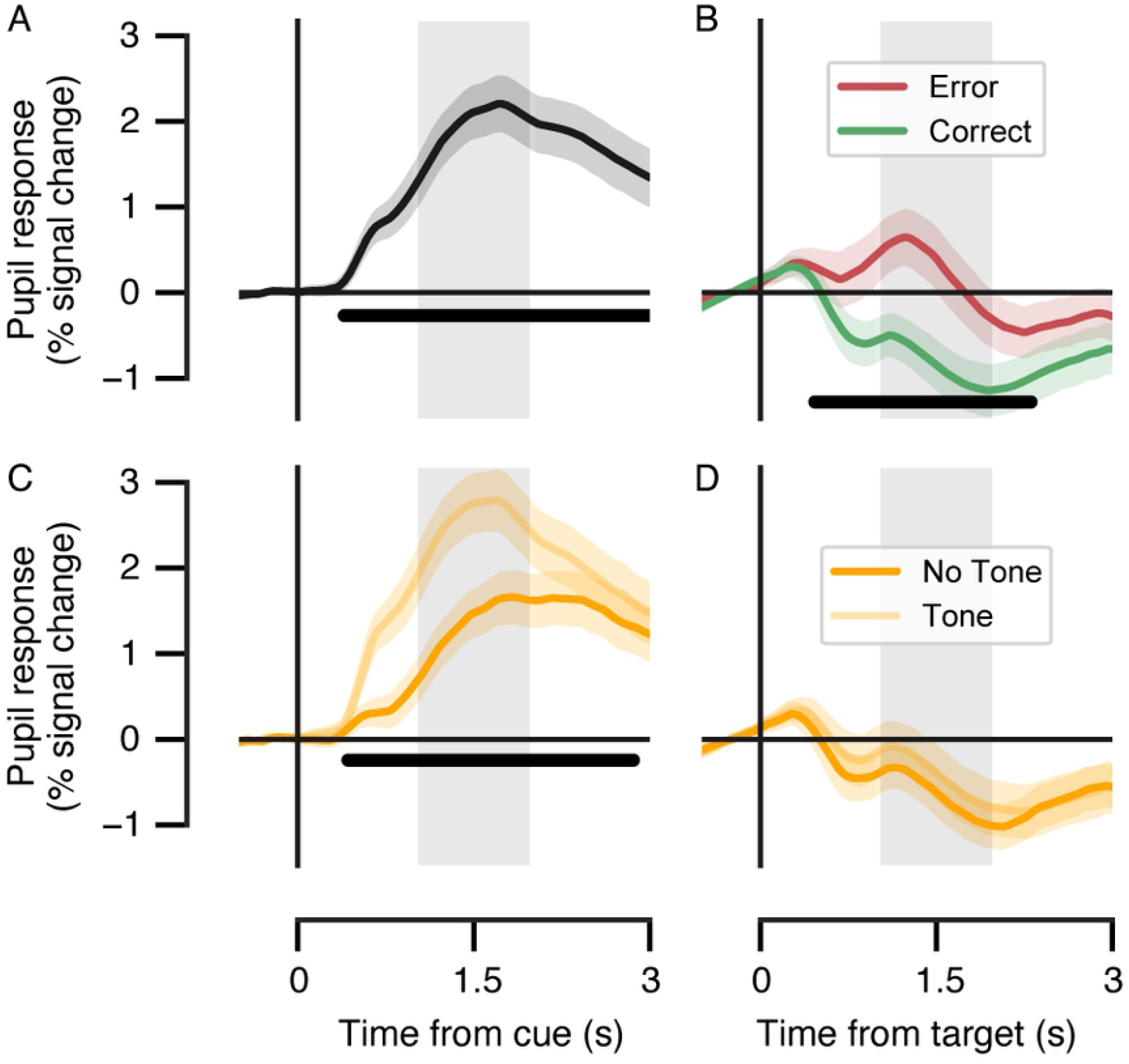
Evoked pupil responses locked to the cue and target in the prediction task across both phases. All trials within the updating and revision phases of the prediction task were included in the evoked pupil response analysis. (**A**) Mean cue-locked pupil responses in the prediction interval. Black bar indicates main effect of cue, *p* < 0.05 (cluster-based permutation test). (**B**) Evoked pupil responses for correct and error trials in the feedback interval (target-locked). Black bar indicates correct vs. error effect, *p* < 0.05 (cluster-based permutation test). (**C**) Evoked pupil responses for tone and no tone trials in the prediction interval (cue-locked) and (**D**) in the feedback interval (target-locked). Black bar indicates tone vs. no tone effect, *p* < 0.05 (cluster-based permutation test). In all panels: variability around the mean responses is illustrated as the 68% confidence interval (bootstrapped; *N* = 30); the grey box indicates the temporal window of interest (1–2 s) with respect to event onset for the phasic pupil responses. Note that the temporal window of interest was independently defined based on the pupil’s canonical impulse response function.

#### Curve fits on target-locked pupil data

Finally, we tested the main hypothesis concerning prediction error dynamics reflected in pupil dilation across the updating and revision phases. Sigmoid curves were fit for each participant’s target-locked pupil response for the tone trials in the high-frequency condition only, separately per phase (updating vs. revision). Individual curve fits are shown in Supplementary Fig 2. Our main hypothesis was that σ would be larger for the high-frequency tone trials in the updating phase as compared with the high-frequency tone trials in the revision phase (i.e., the cue-target mappings were flipped for tone trials only in the revision phase). For the trials without an auditory cue, we did not have expectations about the difference in σ between the updating and revision phases of the task. Therefore, we tested only tone trials for phase-dependent differences (updating vs. revision) of the mean σ parameter (Fig 4B). To examine whether the σ parameters differed, we used Bayesian Wilcoxon Signed-Rank Test (Fig 4B, right column). Whilst differences between σ parameters were obtained, they were not in accordance with our expectations, since σ in the revision phase was larger as compared with the updating phase.

**Figure 4.**
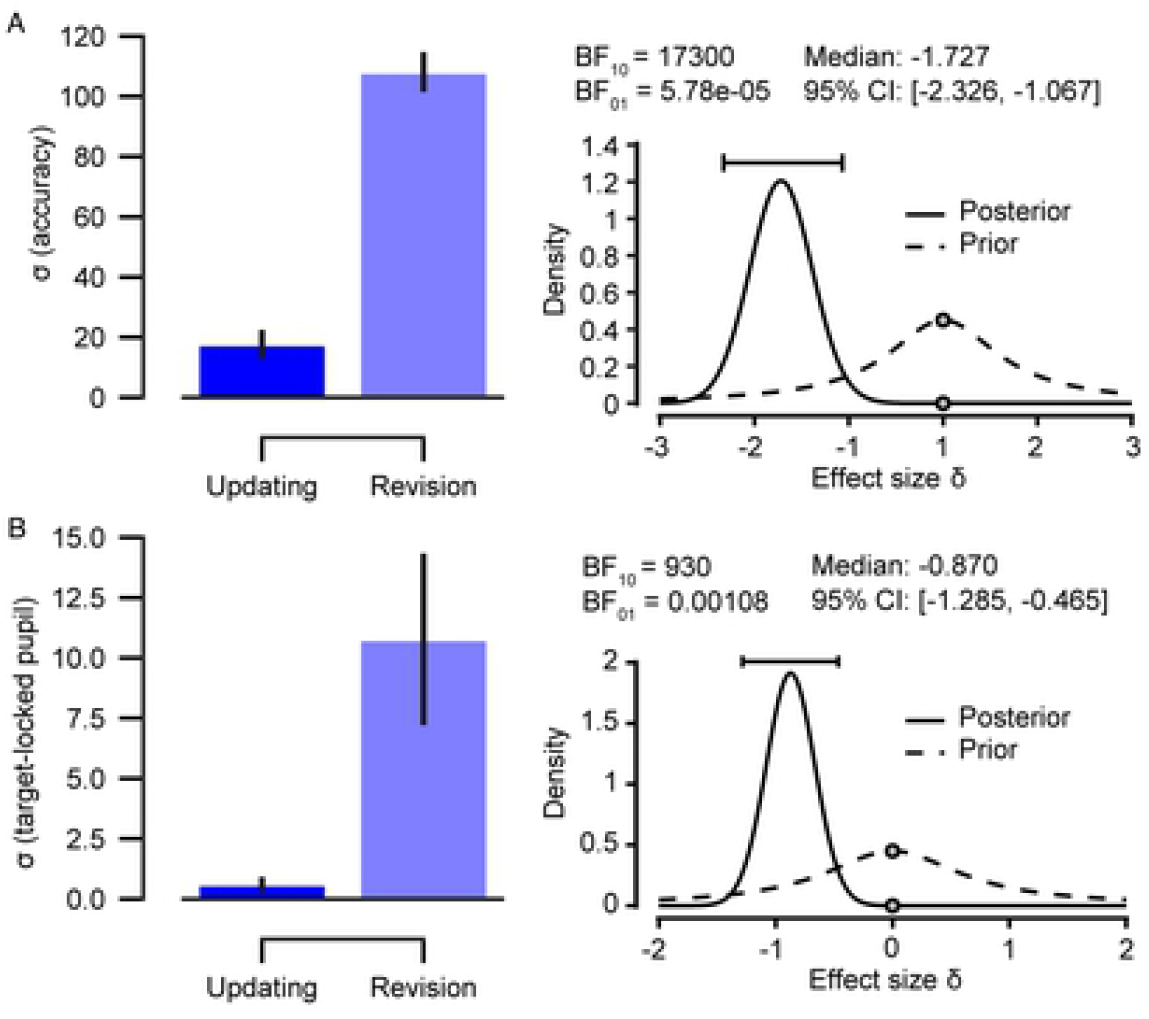
Comparisons of σ parameter of the curve fits on tone trials. For each participant, sigmoid curves were fit to the tone trials (i.e., trials with an auditory cue) and compared between the updating and revision phases of the experiment. (**A)** For accuracy data, all tone trials (i.e., the high- and low-frequency conditions) were used to fit the curves. (**B**) For the target-locked pupil response data, curves were fit to the high-frequency (80%) tone trials only. Note that the cue-target mappings (M1 and M2) which correspond to the high-frequency trials differ per phase depending on the presence of the auditory cue. Results of the Bayesian Wilcoxon Signed-Rank Test are shown for the accuracy and pupil data (right column). Error bars, s.e.m. (*N* = 30).

We also expected that the size of the prediction errors as measured with pupil dilation would decrease as a result of learning, reflected in a negative gain parameter (G) in our curve fits for both phases. In the updating phase, G was negative on average as expected (*M* = −4.1, *SD* = 11.94), however, G was positive on average in the revision phase (*M* = 2.4, *SD* = 9.3), against our hypothesis of reduced prediction errors in both phases of the experiment. Furthermore, it was apparent that the sign of G was not consistent at the individual level (see Supplementary Fig 2).

### Exploratory analyses

Given that the sizes of the σ parameters obtained from the curve fits of pupil data were the opposite of what we had hypothesised, we decided to run an exploratory analysis on the behavioural data. We expected that these additional analyses would provide us with insight into what was going on during the task and would allow us to draw stronger conclusions.

#### Psychometric curve fits on accuracy data

For accuracy data, psychometric curves were fit for each participant’s response accuracy for the tone trials only, separately per phase (updating vs. revision). Individual curve fits are shown in Supplementary Fig 3. As with pupil, our exploratory hypothesis was that σ would be larger for the tone trials in the updating phase as compared with the tone trials in the revision phase (i.e., the cue-target mappings were flipped for tone trials only in the revision phase). For the trials without an auditory cue, we did not have expectations about the difference in σ between the updating and revision phases of the task. Therefore, we tested only tone trials for phase-dependent differences (updating vs. revision) of the mean σ parameter (Fig 4A). To examine whether the σ parameters differed, we used a Bayesian Wilcoxon Signed-Rank Test (Fig 4A, right column). Whilst differences between the σ parameters were obtained, they were not in accordance with our expectations, since σ in the revision phase was larger than in the updating phase.

## Discussion

Our research was motivated by the observation that learning in the predictive-processing framework is often reduced to Bayesian model updating (Perfors, 2012; Schulz, 2012; Gentner & Hoyos, 2017). We argued that existing accounts of learning in the predictive-processing framework currently lack a crucial component: a constructive learning mechanism that accounts for changing models structurally when new hypotheses need to be learnt. Kwisthout and colleagues (2017) proposed that model revision is a learning mechanism that is distinct from Bayesian model updating and accounts for such a structural change in generative models. Similarly, Heald and colleagues (2022) have recently presented a theory for sensorimotor learning, called contextual inference, that differentiates between the adaptation of behaviour based on updating of existing and creation of new motor memories and adaptation due to changes in the relative weighting of these motor memories. Building on this, we investigated whether we could empirically distinguish between Bayesian model updating and model revision. We expected to be able to differentiate between the two mechanisms based on their prediction error dynamics, which we assumed to be reflected in pupil dilation (Preuschoff et al., 2011; Browning et al., 2015; O’Reilly et al., 2013; Colizoli et al., 2018; Lawson et al., 2017). After obtaining unexpected results based on pupil data, we formed additional, exploratory hypotheses based on the prediction dynamics reflected in accuracy. To investigate our question, we created a within-subject computer-based experiment with two phases (the updating and the revision phase). In the updating phase, participants had to learn and predict the probabilistic relationship between the cues and a target stimulus, and in the revision phase, a change occurred to what had been learnt previously.

To test whether participants performed the task as expected, we performed some basic quality controls on behavioural and pupil data. Behavioural data showed that participants on average correctly predicted the cue-target contingencies in both experimental phases. This was reflected in the cue-target mapping effect for tone and no tone trials in the updating phase, and a reversed effect of cue-target mapping effect was observed in the revision phase for tone trials (when the mapping switched) only. As expected, a similar pattern was also observed in pupil data.

Before turning to the main research question, the quality of the pupil measures was checked and compared to effects observed in previous literature. Decision preparation and the preparation of the motor response were reflected in the evoked pupil response to the visual and auditory cue onsets replicating previous work (Colizoli et al., 2018; Urai et al., 2017). We also observed the peak of the group-level cue-locked evoked response around 1.5 seconds, which is in line with previous findings (e.g., Mathôt, 2018). Furthermore, erroneous responses resulted in larger pupil dilation as compared to correct responses, an effect that has also been previously reported (Braem et al., 2015). Finally, we found a well-known auditory effect on pupil responses with pupil responses being bigger on trials with a tone compared to trials without a tone (e.g., Liao et al., 2016).

Given that participants understood and correctly performed the task, we turn to the results related to our main hypotheses. Multiple studies (Friston et al., 2016; Smith et al., 2020; Fitzgerald, 2015; Clark, 2016; Zenon, 2019) have shown that prediction error decreases as a result of learning. Building on this we hypothesised that in the updating and the revision phase, different temporal dynamics in prediction error decrease would be observed. We hypothesised that in the updating phase participants would be gradually learning the probabilistic relationship between the cues and the target orientation, leading to model updating. As a result of model updating, participants should become better at predicting future sensory input. New sensory observations should thus yield prediction errors that are smaller than the preceding ones, and the degree to which the model needs to be updated should decrease over time. In other words, we expected prediction error to decrease *gradually* over time. In the revision phase the rules switched for the tone trials, which should initially lead to an increase in prediction error. When the change in rules is learned, a new hypothesis is added to a learner’s model, resulting in model revision. An integration of a new hypothesis should lead to an *abrupt* decrease in prediction errors (i.e., an “aha” moment). Whilst prediction errors reflect the deviation between expectations and target-feedback presentation, the behavioural responses (i.e., button presses) are supposed to reflect the *predictions* themselves. Thus, we expected to find a similar pattern of responses in accuracy data as in prediction errors but in the opposite direction.

Curve fitting revealed significant differences between the phases of the task in pupil and accuracy data as reflected in the σ parameter. However, the difference was in the opposite direction as expected; the σ was significantly smaller in the updating phase compared to the revision phase, in accuracy and pupil data. These results suggest that the participants were learning more gradually in the revision phase compared to the updating phase of the experiment, contrary to our expectations.

One possible interpretation of these results is that our experimental manipulation induced model revision in the updating phase and model updating in the revision phase. It might have been that in the updating phase participants built multiple internal models, that they thought could capture the structure of the task, from scratch. As they were learning the task they were alternating between these models and upon the realisation of the rule participants settled for the correct model, resulting in a rapid decrease in prediction error and accuracy as the data in the updating phase shows. We assumed we prevented participants from learning models from scratch in the updating phase by providing them with detailed instructions and pictorial representation of the stimuli resented in the task, before the task started. By that, we thought, we equipped participants with a crude model that would contain hypotheses about all the relevant variables of the task. However, in light of the current results we believe that our instructions did not result in the construction of a simple model that participants could use as a baseline upon entering the task.

At the beginning of the revision phase, participants expected for the rules of the task to change. All the experimental variables (e.g., target, visual cues) in the revision phase were the same as in the updating phase, possibly signalling to the participants that one of the models that they had already constructed in the updating phase could be suitable for explaining the change in the revision phase. If this was the case, then participants in the revision phase merely reused a correct, existing model, and started gradually updating that model rather than adding a new hypothesis (based on a new experimental variable) to a model they had built in the updating phase.

We could possibly have avoided participants building models from scratch in the updating phase had we ran a computerised familiarisation phase with all the relevant experimental variables with the participants prior to the experiment. This would make sure that participants have constructed crude models of the task before the experiment started and that they could then use and update in the update phase. Additionally, to make sure that participants in the revision phase were not just reusing one of the models they had constructed in the updating phase but instead build on an existing model, we could have introduced a new experimental variable in the revision phase that was present in the familiarisation but not in the updating phase. In that case, participants would have to add a new hypothesis to a model (instantiating model revision) constructed in the updating phase, if they were to successfully learn the new rule in the revision phase.

Our results also revealed that the gain parameter, G, which indicates the direction of the σ parameter, was negative in the updating phase as expected and positive in the revision phase contrary to our expectations. This suggests that pupil dilation and hence prediction error was on average decreasing in the updating phase and increasing in the revision phase. These results are unexpected, because, according to existing literature, prediction error is decreasing as a result of learning (Friston et al., 2016; Smith et al., 2020; Fitzgerald, 2015; Clark, 2016; Zenon, 2019) and so should pupil dilation (Kayhan, 2019). However, a look at the individual level pupil data (see supplementary Fig 2) reveals large variability in the sign of the σ parameter for both phases, potentially suggesting individual differences in the size of pupil dilation over time. This suggestion is in line with recent findings of substantial inter- and intra-individual variation in the size of pupil dilation over trials (Sybley et al., 2020). More specifically, the study shows that in a simple digit-span memory task pupil dilation was consistently increasing over trials for some participants and for others it was decreasing. There were also participants for whom the trend was changing throughout the task. Another factor that could explain why pupil dilation was increasing in the revision phase is that participants in this phase were more fatigued than in the updating phase at the beginning of the experiment. As a consequence, they had to exert more effort to maintain concentrated on the task and process the change in the task rules. Increased cognitive effort would result in increasing pupil dilation as has been shown many times before (for review see van der Wel & van Steenbergen, 2018; Hyönä et al., 1995; Porter et al., 2007).

All in all, our data shows that there exists a qualitative difference between model updating and model revision, following the initial theoretical proposal from Kwisthout and colleagues (2017). However, model updating seemed to have occurred in the model revision phase and model revision in the model updating phase of the experiment for reasons described above. Future studies should therefore make sure to induce experimental manipulations that have the initially intended effect. Alternatively, future studies might investigate empirically the interpretation of the current results: that participants construct multiple models at the beginning and later choose among them and update them. Lastly, our study is one of the few that studied how the change in prediction error, as captured by pupil dilation, over time (for a few exceptions see Kayhan, 2019; Koenig et al., 2017). Therefore, little is known about how pupil dynamics and hence prediction errors change over extended periods of time and whether individual differences exist in this process. More studies should thus examine pupil dilation and hence prediction error in such a manner in the future.

## Supplementary material

**Supplementary Figure 1.**
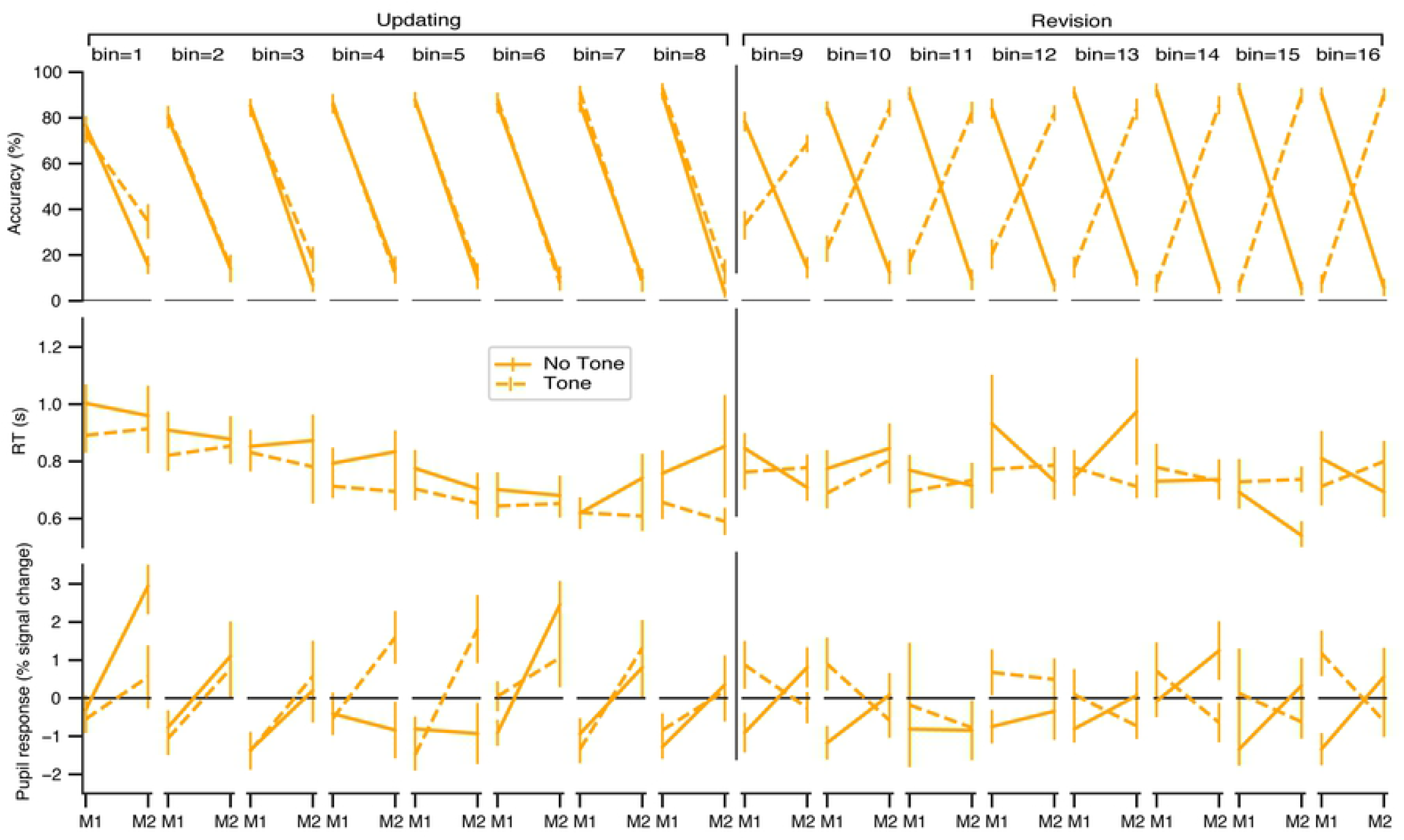
Cue-target prediction task results as a function of trial bin. Prediction accuracy (top row), mean RT (middle row), and target-locked pupil dilation (bottom row) as a function of visual cue-target mapping condition (M1 vs. M2), the presence of the auditory cue (tone vs. no tone), and consecutive trial bin. Trial bins 1-8 correspond to the updating phase, and trial bins 9-16 correspond to the revision phase (25 trials per bin).

**Supplementary Figure 2.**
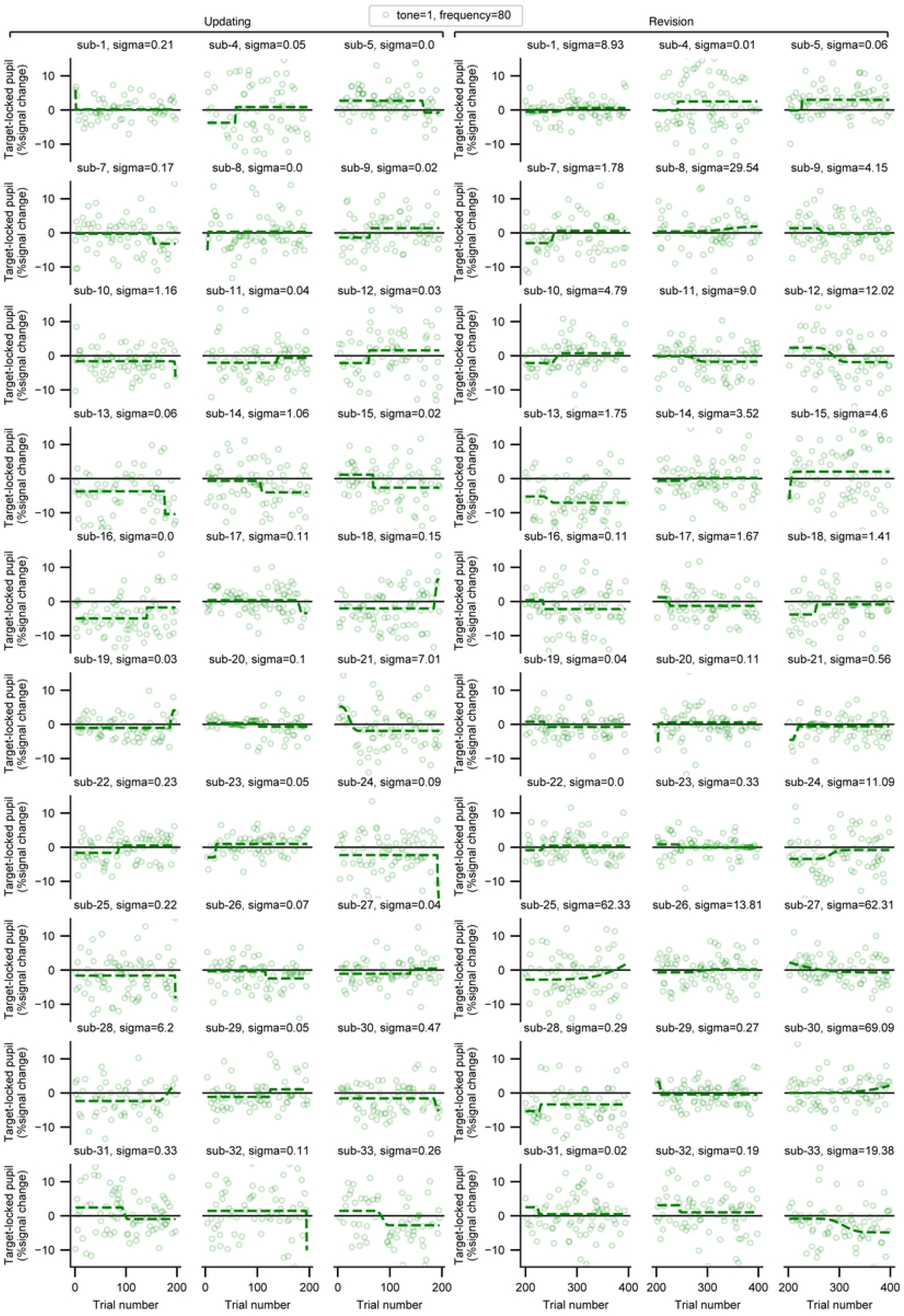
Curve fits for the target-locked pupil response (single-trial). Individual curves were fit on the target-locked pupil response data for the high-frequency tone trials (i.e., the auditory cue and 80% frequency condition) separately for each phase (updating vs. revision).

**Supplementary Figure 3.**
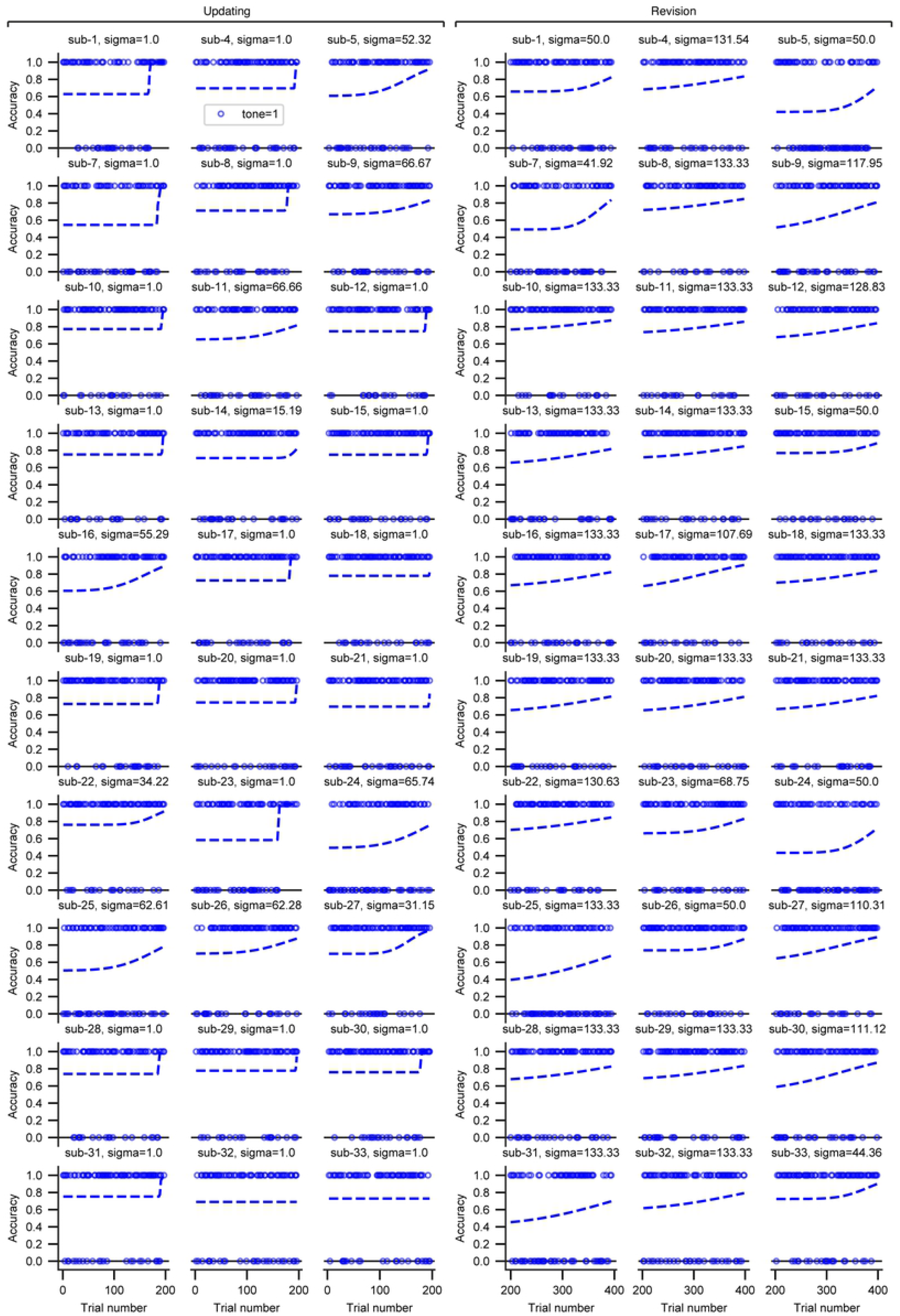
Psychometric curve fits for accuracy (single trial). Individual psychometric curves were fit on the response accuracy data for the tone trials (auditory cue condition) separately for each phase (updating vs. revision).

**Supplementary Table 1.** Descriptive statistics on free parameters from curve fits for the accuracy and pupil data.

| | $\mu$ | | $\sigma$ | | $a0$ | |
| --- | --- | --- | --- | --- | --- | --- |
|  | Updating | Revision | Updating | Revision | Updating | Revision |
| *Valid* | 30 | 30 | 30 | 30 | 30 | 30 |
| *Missing* | 0 | 0 | 0 | 0 | 0 | 0 |
| *Median* | 191.29 | 400 | 1 | 132.437 | 0.711 | 0.649 |
| *Mean* | 187.996 | 389.102 | 17.737 | 108.104 | 0.69 | 0.617 |
| *Std. Deviation* | 13.053 | 22.626 | 26.1 | 36.207 | 0.08 | 0.108 |
| *Minimum* | 156.964 | 323.083 | 1 | 41.918 | 0.492 | 0.351 |
| *Maximum* | 200 | 400 | 66.667 | 133.333 | 0.779 | 0.77 |

*Pupil response*
| | $\mu$ | | $\sigma$ | | $a0$ | | $G$ | |
| --- | --- | --- | --- | --- | --- | --- | --- | --- |
|  | Updating | Revision | Updating | Revision | Updating | Revision | Updating | Revision |
| *Valid* | 30 | 30 | 30 | 30 | 30 | 30 | 30 | 30 |
| *Missing* | 0 | 0 | 0 | 0 | 0 | 0 | 0 | 0 |
| *Median* | 130.827 | 246.795 | 0.099 | 1.766 | -1.269 | -0.365 | -2.522 | 1.036 |
| *Mean* | 122.354 | 262.328 | 0.612 | 10.772 | 0.767 | -1.99 | -4.136 | 2.397 |
| *Std. Deviation* | 63.868 | 60.348 | 1.656 | 19.487 | 9.846 | 8.267 | 11.938 | 9.307 |
| *Minimum* | 0.747 | 186.918 | 1.369e -5 | 8.487e -4 | -5.082 | -43.248 | -51.324 | -7.209 |
| *Maximum* | 198.993 | 457.277 | 7.008 | 69.092 | 51.521 | 6.503 | 8.48 | 45.274 |

### Prediction error in pupil dilation explored as the 4-way interaction between accuracy, cue-target mapping, auditory cue, and phase

If the full scope of the prediction error space was measurable in the target-locked pupil dilation, a 4-way interaction between the factors: accuracy (error vs. correct), and cue-target mapping (M1 vs. M2), auditory cue (tone vs. no tone), and phase (updating vs. revision). For instance, a *correct* response on a *low frequency* (20%) trial may reflect a wrong button press on the part of the participant but will elicit a substantial prediction error due to the unexpected outcome once the contingencies are learned. The size of the two-way interaction term defined by the factors accuracy and cue-target mapping should change over time in accordance with the auditory-cue rules in the updating and revision phases, representing the deviation between expectation and outcome in the form of the target orientation.

However, we did not proceed with analysis due to too many missing cases across the 16 conditions determined by the 4-way interaction (max. 14/30). These missing cases were due to the rare occurrence of certain conditions, such as correct and low frequency trials. Therefore, we could not make any inference on the potential dynamics of this 4-way interaction in the current task design.

